# DeepUnitMatch: tracking neurons across days in electrophysiology using Deep Neural Networks

**DOI:** 10.64898/2026.01.30.702777

**Authors:** Suyash Agarwal, Wentao Qiu, Kenneth D Harris, Enny van Beest, Célian Bimbard

**Author notes:** These authors contributed equally (Suyash Agarwal, Wentao Qiu). These authors contributed equally (Enny van Beest, Célian Bimbard).

## Abstract

To understand neural processes such as learning or memory, we need to track the activity of populations of neurons at the level of single spikes and across days. Here, we leverage deep neural networks to build DeepUnitMatch, a software that reliably tracks individual neurons in high-density electrophysiological recordings across weeks. DeepUnitMatch uses only the spike waveforms of the neurons, and not their spiking patterns, and outperforms current solutions.

## Introduction

Tracking the millisecond, single-spike activity of large populations of neurons across days is fundamental for understanding the neural basis of many cognitive processes, such as learning and memory. Calcium imaging methods allow long-term tracking of neuronal populations in cortex^1–5^ but lack the temporal precision to capture fast dynamics. By contrast, high-density extracellular electrophysiological recordings with Neuropixels probes enable stable, chronic measurements of hundreds of neurons with high spatiotemporal resolution across the brain^6–12^.

A central challenge in electrophysiology is to track individual neurons across recordings. Several methods have been proposed that do not rely on functional properties of the neurons for matching, and are thus suited for characterizing changes in the neurons’ response properties. The first method relies on concatenating pairs of recordings and spike-sorting the resulting file^9^. However, it does not scale to the multi-session recordings common in modern experiments. Recent methods, such as EMD^13^ and UnitMatch^14^, address these limitations by spike sorting each session independently and then registering units across recordings using engineered waveform features. While more effective and accurate than the concatenation method^14^, these approaches still have missed or false matches, potentially because they depend on predefined human-engineered features that fail to capture the full complexity of neuronal waveforms.

Deep learning offers a powerful, unsupervised alternative. Neural networks have been successfully used to track neurons in calcium imaging^5^, as well as to perform tasks in electrophysiology such as cell-type classification^15–17^, spike-sorting^18^ and merging split spike clusters^19^. This suggests that they may be particularly suited for extracting rich and relevant features from the neurons’ waveforms.

Here, we describe a new method, DeepUnitMatch, that uses deep neural networks to identify neuronal spike waveforms across multiple recording sessions. DeepUnitMatch outperforms the current best method, UnitMatch, because 1) it finds more matches across days and 2) these matches are better aligned with functional matching scores, indicating better performance.

## Results

DeepUnitMatch uses the same pipeline architecture as UnitMatch (Figure 1a-c) but with an unsupervised, deep neural network based method for waveform feature extraction (Figure 1b, steps 1-3). To generate the probability of each pair of recorded neurons being a match, DeepUnitMatch uses a deep convolutional neural network to compute similarity scores between pairs of neurons (Figure 1b), instead of the engineered similarity features of UnitMatch.

**Figure 1.**
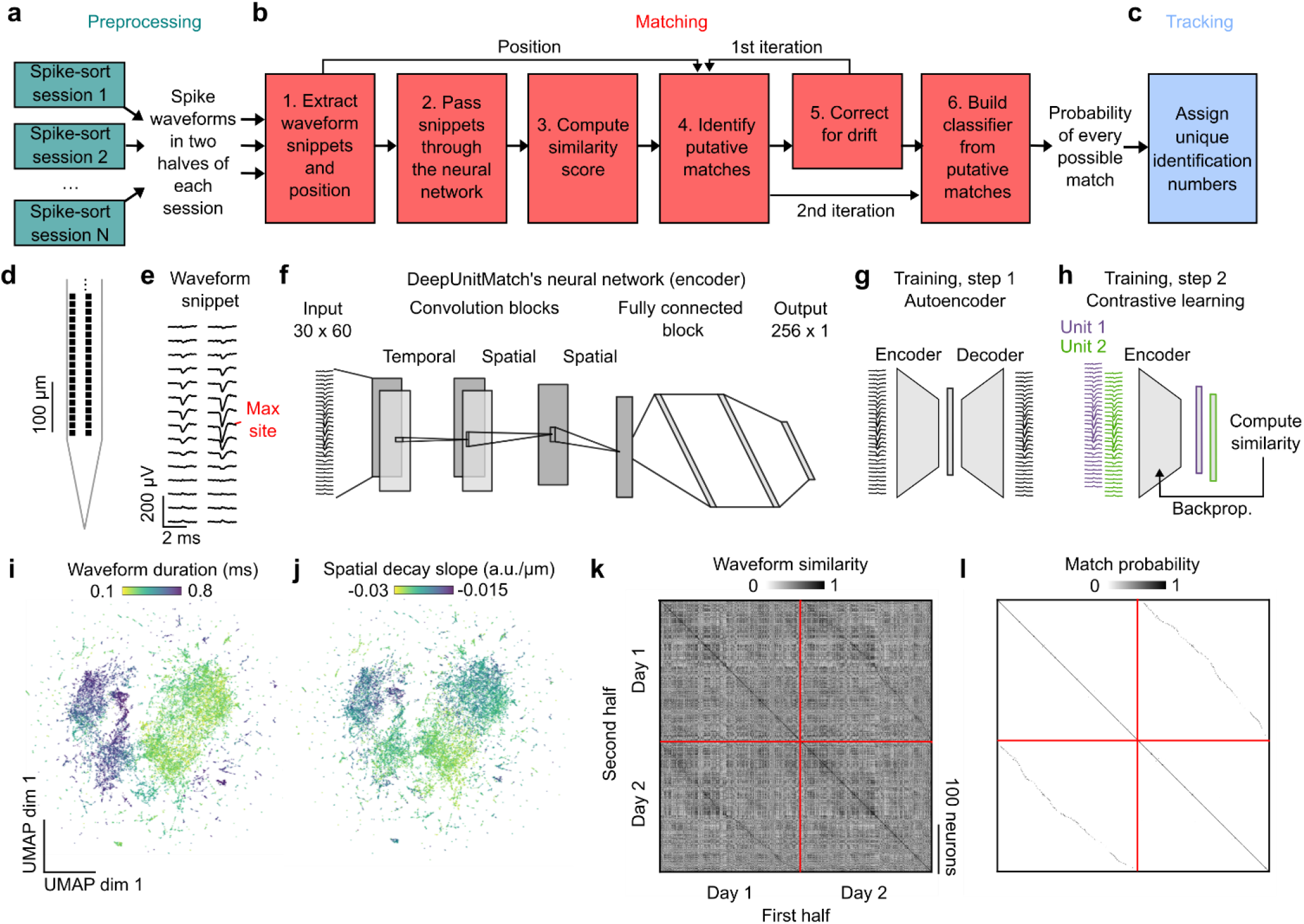
DeepUnitMatch pipeline. **a**, Preprocessing steps. **b**, Matching steps. **c**, Tracking step. **d**, Schematics of a Neuropixels 2.0 probe, where black squares correspond to recording sites. **e**, Example waveform snippet, across 30 channels. **f**, Architecture of DeepUnitMatch’s deep neural network (encoder). **g**, First step of the encoder’s training, to reconstruct individual waveforms with an autoencoder. The decoder has a mirrored architecture **h**, Second step of the encoder’s training, to give similar (versus different) outputs for waveforms from the same (versus different) neurons, with contrastive learning. **i**, 2-D UMAP plot of neural network representations for each waveform, colored by waveform duration (measured from peak to trough, in µs). Neurons across all sessions and animals are shown. **j**, Same as **i**, colored by spatial decay slope, which measures how quickly the waveform decays from its maximum peak along the electrode channels. **k**, Similarity matrix of the neurons’ waveforms for two example sessions. The main diagonal compares the two copies of a neuron’s waveform from two halves of a recording session, while the two offset diagonals show highly similar waveforms across sessions. Neurons are sorted by depth on the probe within days. **l**, Same as **k**, with the match probability matrix for the same two sessions.

As with UnitMatch, DeepUnitMatch takes as an input the spike sorted waveforms of individual recording sessions. We focused on Neuropixels 2.0 recordings^9^ (Figure 1d), used Kilosort to spike-sort them^20,21^, and extracted the spatiotemporal spike waveforms from the first and second half of the recording for each (putative) neuron, defined as a well-isolated single unit^22^ (Figure 1a). For each neuron, we found the channel with the maximum amplitude and defined a ‘waveform snippet’ including the 30 closest channels to this maximal channel (Figure 1e). The waveform snippets are the input to the DeepUnitMatch neural network architecture, which is thus oblivious to the neurons’ spatial position on the probe.

DeepUnitMatch then extracts a low-dimensional representation of each neuron’s waveform using a deep convolutional neural network. The network consists of temporal and spatial convolutional blocks (each containing several layers) and a fully connected block to form a latent representation of each waveform (Figure 1f), reducing the size of the representation from 1800 to 256 parameters. The network is trained in two steps. First, to extract general representations of spike waveforms, we plugged the network into an autoencoder^23^ tasked with reconstructing the original waveforms from the latent representation (similarly to Ref.^16^, Figure 1g). Next, we further trained the network to learn different representations for waveforms from different neurons and similar representations for waveforms from the same neuron (Figure 1h). To do so, we used contrastive learning^24^ on the waveform snippets from the two halves of a recording for neurons in individual sessions. The small differences in waveforms between the two halves act as a natural form of augmentation. In the end, the waveform representations at the output of the encoder network captured known spike waveform features such as spike width (Figure 1i) and spatial decay (Figure 1j), which typically form an electrical signature specific to each neuron.

DeepUnitMatch finally computes the waveform-based similarity across all pairs of neurons to track individual neurons across days. DeepUnitMatch first computes the cosine similarity between the latent representations of all neurons across all recording sessions that are given by the user, yielding a global pairwise similarity matrix (Figure 1k).

DeepUnitMatch then combines this waveform similarity matrix with the matrix of the pairwise Euclidean distance between neurons to select putative matches. Similarly to UnitMatch, DeepUnitMatch uses these putative matches to build a Naive Bayes classifier and obtain a posterior probability of each pair of neurons across the recordings being a match (Figure 1l). This posterior probability can then be used to assign unique identities to individual neurons across all recordings^14^.

To evaluate the matching performance, we used the neurons’ functional features, which are not used for matching. Ground-truth data of neurons tracked across days in electrophysiology recordings does not exist. However, some functional properties of the neurons can be used as independent fingerprints to assess the algorithms’ performance^9,14,25–40^. For example, inter-spike interval (ISI) distributions are reliable within neurons (Figure 2a) yet different across neurons (Figure 2b), and are stable across days (Figure 2c)^14,25,28,40^. Therefore, we can use the correlation of ISI-distributions to independently assess whether a pair of neurons is likely to be different (low correlation) or the same (high correlation) (Figure 2d). Using these correlations, we computed the area under the receiver operating curve (AUC) which discriminates between matches and non-matches across days. Higher AUC values indicate better algorithmic matching performance. This measure of performance is highly sensitive to false positives, and much less to false negatives (Supplementary Figure 1).

**Figure 2.**
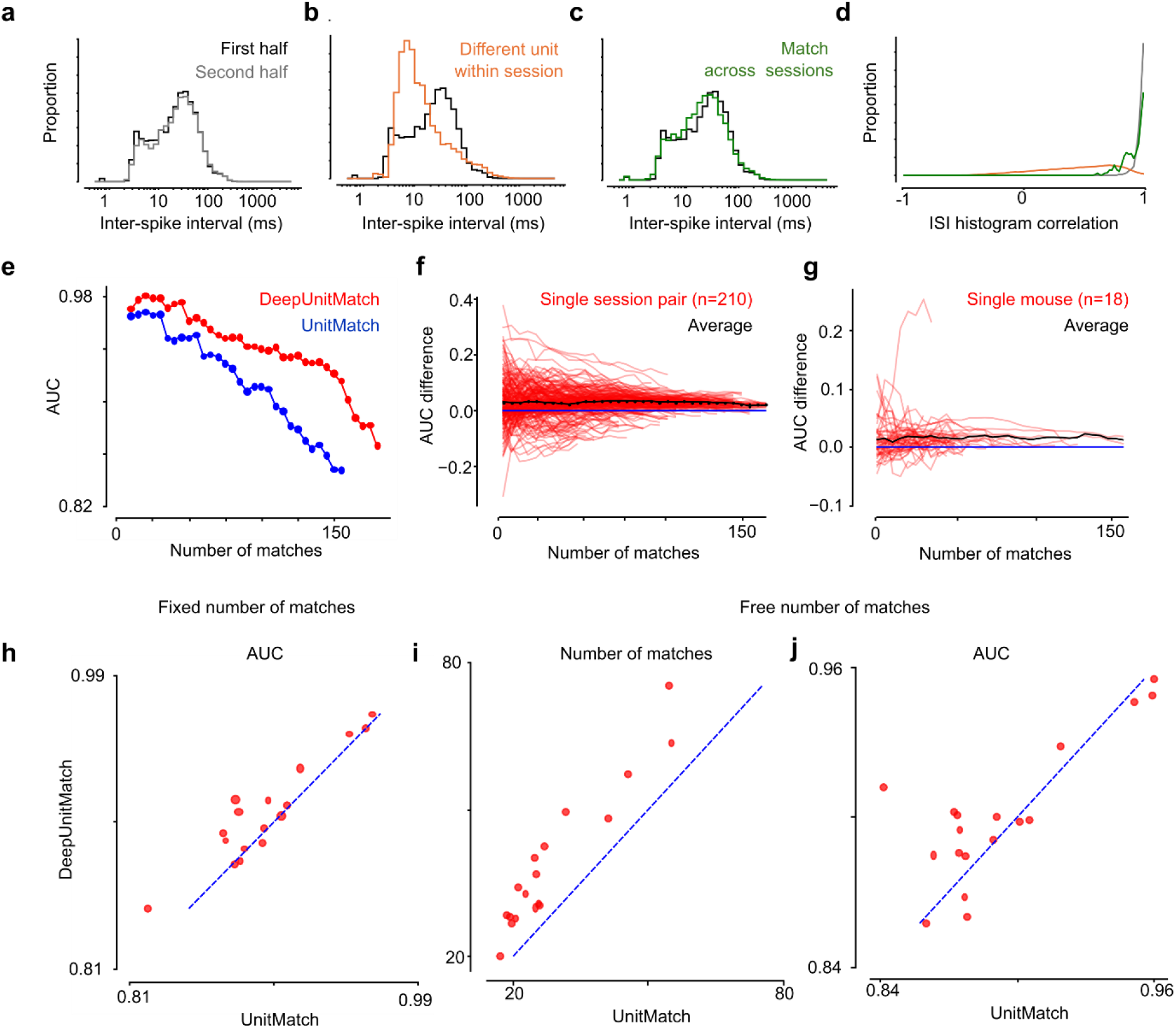
DeepUnitMatch outperforms current best algorithm UnitMatch. **a**, Histograms of inter-spike intervals (ISIs) for an example neuron in the first (*black*) and second (*grey*) half of a session. **b**, Same as **a**, with the histogram of a different unit within the same session (*orange*). **c**, Same as **a**, with the histogram of the same unit tracked across another session (*green*). **d**, Distribution of the ISI histogram correlations for two copies of the same unit (*grey)*, different units from the same session (*orange*), and matches across sessions (*green*). **e**, AUC values when increasing the number of matches selected for DeepUnitMatch (*red*) and UnitMatch (*blue*). **f**, Difference in AUC between DeepUnitMatch and UnitMatch, with one line per pair of recordings (*red*) and their average (*black*) for an example animal. This animal had 21 recording sessions, so there are 210 session pairs. **g**, Same as **f** but each line is the average AUC difference for all pairs of recordings from a given mouse (*red*), and averaged over 18 mice (*black*). **h**, Average AUC of DeepUnitMatch plotted against UnitMatch, one dot per mouse. The number of matches returned by both algorithms was determined by the number found by UnitMatch. The size of each point is proportional to the average number of matches found across recordings from that mouse. **i**, Same as **h**, for the number of matches when the algorithms have their default threshold (free number of matches). **j**, Same as **h** but each algorithm sets its own number of matches, as shown in **i**.

DeepUnitMatch is more accurate than UnitMatch (Figure 2e-h). The AUC value depends on the number of matches selected, as exemplified in an example session (Figure 2e): a simple change in selection threshold (conservative vs. liberal) can have a drastic effect on the algorithm’s performance. To ensure a fair comparison between algorithms, we computed the AUC values across all possible numbers of selected matches. In this example pair of sessions, DeepUnitMatch outperformed UnitMatch for any given number of matches. This was also true on average across all possible pairs of sessions for this example mouse (Figure 2f), and across all 18 mice (Figure 2g). To statistically quantify this increase in performance, we fixed the number of matches to that found by UnitMatch. While this strategy should be to the advantage of UnitMatch, DeepUnitMatch consistently outperformed UnitMatch (p = 0.007, n = 17 mice, paired Wilcoxon signed-rank test, Figure 2h; one mouse did not yield enough matches across sessions), suggesting overall a higher accuracy.

The different stages of training were useful to make DeepUnitMatch more accurate. We used the same approach to quantify the performance of different models that had been only partially trained. While a completely untrained model still achieved reasonable performance thanks to spatial and suboptimal waveform information, the fine-tuning with contrastive learning and the auto-encoder training improved performance to reach its highest level in the fully trained model (p = 4.0 x 10^-7^, n = 18 mice, Friedman test; DeepUnitMatch vs DeepUnitMatch untrained : p = 3.1 x 10^-5^, DeepUnitMatch vs DeepUnitMatch no fine-tuning : p = 6.6 x 10^-4^, DeepUnitMatch vs DeepUnitMatch no AE tuning : p = 9.3 x 10^-3^, paired Wilcoxon signed-rank tests, Supplementary Figure 2).

When letting each algorithm decide on the number of selected matches, DeepUnitMatch found ∼40% more matches than UnitMatch. This was true in all 18 mice (38 ± 4% more matches, mean ± s.e., p = 7.6 x 10^-6^, n = 18 mice, paired Wilcoxon signed-rank test, Figure 2i), while both algorithms yielded similar accuracies (p = 0.15, n = 18 mice, paired Wilcoxon signed-rank test, Figure 2j). Importantly, since our performance measure particularly penalizes algorithms with high false positives rates (Supplementary Figure 1), our results suggest that the additional matches found by DeepUnitMatch were real matches missed by UnitMatch.

DeepUnitMatch outperforms UnitMatch when tracking neurons across many sessions. Since matching performance of UnitMatch drops with number of days in between recording sessions^14^, we wanted to test whether DeepUnitMatch maintained a better matching performance over longer periods of time. We first compared the matching performance of UnitMatch and DeepUnitMatch with their respective default parameters as a function of the number of days between pairs of recordings (Δ days). In the example mouse, the number of matches found by DeepUnitMatch was higher than that found by UnitMatch across all Δ days, and these matches were more accurate, even after >100 days (Figure 3a,b). This was generally true across all mice (Figure 3c,d). Beyond matching neurons across pairs of sessions, the original UnitMatch algorithm also contains a module for tracking individual neurons across multiple sessions. We thus ran the UnitMatch tracking algorithm on the probability outputs from both UnitMatch and DeepUnitMatch. The probability of single neurons being tracked across recordings was higher for DeepUnitMatch than UnitMatch, and it was done more accurately (Figure 3e,f; proportion of tracked neurons depends on interaction between number of days and algorithm: F_14,404_ = 1.95, p = 0.02; AUC depends on algorithm F_1,386_ = 13.0, p = 0.004, and the interaction with number of days F_14,386_ = 1.93, p = 0.02).

**Figure 3.**
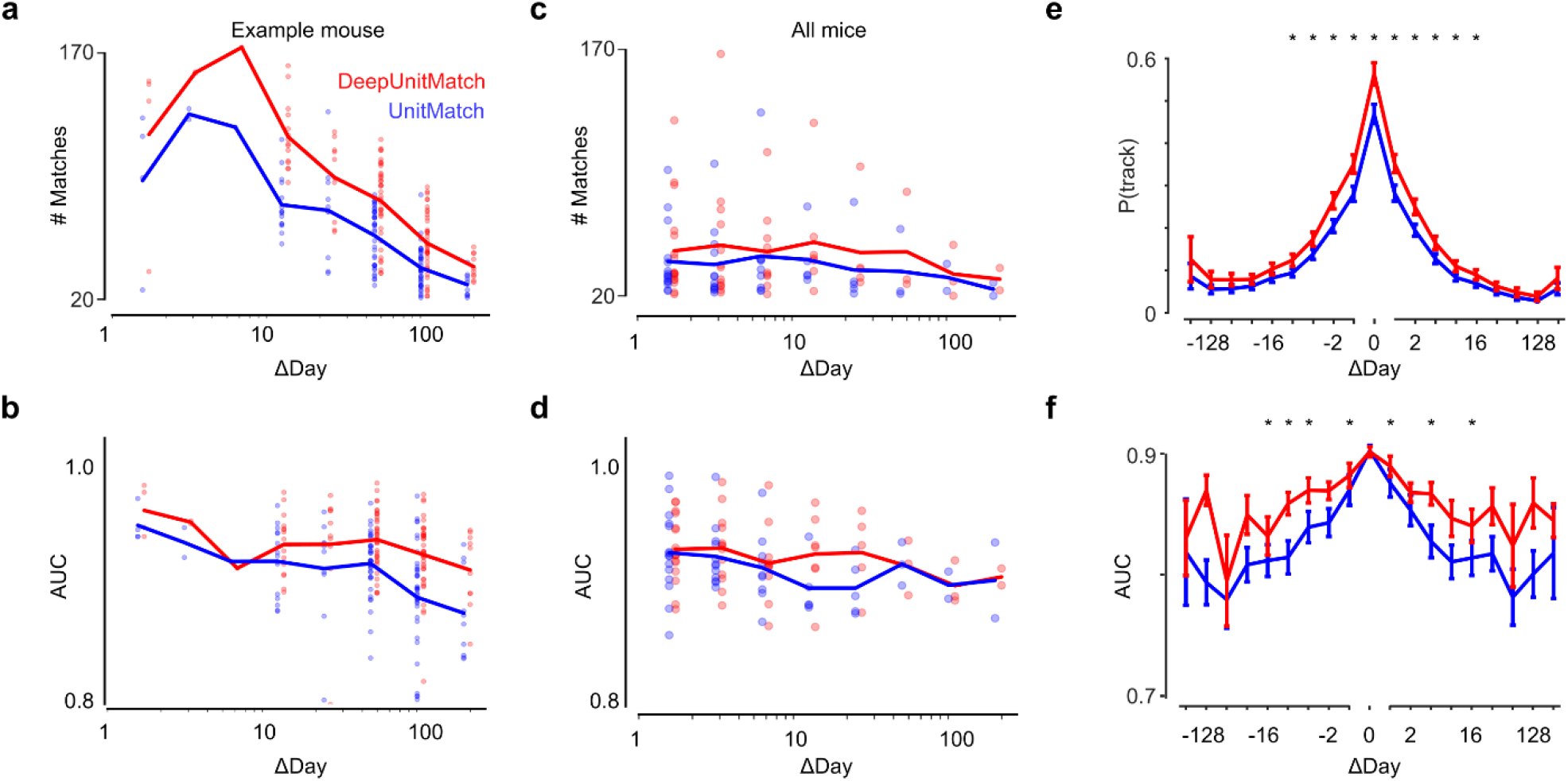
DeepUnitMatch tracks more neurons than UnitMatch, and more reliably, across days and weeks. **a**, Number of matches found by DeepUnitMatch (*red*) and UnitMatch (*blue*) for pairs of recordings with varying number of days in between, for all recordings from a single mouse (*thick line*, average across sessions). **b**, Same as **a**, with the AUC values. **c**, Same as **a**, for 18 mice (*thick line*, average across mice). **d**, Extending **b** to 18 mice. **e**, The probability of tracking neurons using DeepUnitMatch and UnitMatch, as a function of the interval (measured in days) between any pair of recording sessions. **f**, Same as e, with the average AUC of each method plotted. * indicates p < 0.05 difference between DeepUnitMatch and UnitMatch for a specific bin. Paired non-parametric tests after significant main effect.

## Discussion

Tracking the activity of neurons in electrophysiology has remained a long-standing challenge. Waveform-based algorithms are usually complemented with functional measures to track individual neurons^29– 32,37–39,41–45^. This may create circularity when the underlying scientific question is about the stability of the neurons’ functional properties. It is thus crucial to build algorithms that use only the neurons’ waveform to track them across recordings. The current best method UnitMatch^14^ uses human-engineered waveform features. Here, we show that deep neural networks can extract richer and more reliable features from the waveforms, outperforming UnitMatch both in number of matches and accuracy, even with many days between recordings.

Our waveform feature extraction module is similar to those recently developed for other waveform-based tasks^15–17^. The advance of more complex networks, and larger dataset bases, may help in creating even more efficient networks for these various waveform-based tasks. Importantly, this deep neural network-based approach will also benefit from the improvement of spike sorting algorithms^21^, but also of clustering algorithms^45^, which we did not address here.

By making DeepUnitMatch part of the already efficient UnitMatch pipeline available via GitHub (https://github.com/EnnyvanBeest/UnitMatch/tree/main/UnitMatchPy) and pypi.org (https://pypi.org/project/UnitMatchPy/), which in turn can be integrated with SpikeInterface^46^, we minimized the steps needed from users to implement this change. Indeed, our new algorithm is readily applicable and does not require changes in the waveforms’ files format. It can also be applied to merge units that were mistakenly split by the spike-sorting algorithm^19^.

## Acknowledgements

We thank C. B. Reddy for lab management and the Cortexlab for useful discussions and feedback. This research was funded by the Wellcome Trust (Investigator Award 223144/Z/21/Z to K.D.H., and Early Career Award 227065 to C.B.) and the European Union’s Horizon 2020 (Marie Skłodowska-Curie grant agreement no. 101022757 to E.H.v.B.)

## Author contributions

Conceptualization: E.H.v.B. and C.B.; methodology: S.A., W.Q., E.H.v.B. and C.B.; software: S.A. and W.Q.; formal analysis: S.A. and W.Q.; resources: E.H.v.B. and C.B.; writing—original draft: S.A., W.Q., E.H.v.B. and C.B.; writing—review and editing: S.A., W.Q., K.D.H., E.H.v.B. and C.B.; visualization: S.A., W.Q., E.H.v.B. and C.B.; supervision: E.H.v.B. and C.B.; funding acquisition: K.D.H., E.H.v.B. and C.B.

## Data availability

Part of the data is already available online^14^ (https://doi.org/10.6084/m9.figshare.24305758.v1, https://doi.org/10.5522/04/24411841.v1), and the rest will be made available upon publication.

## Code availability

The code is available on UnitMatch’s GitHub (https://github.com/EnnyvanBeest/UnitMatch/tree/main/UnitMatchPy).

## Methods

Experimental procedures were conducted at UCL according to the UK Animals Scientific Procedures Act (1986) and under personal and project licenses released by the Home Office following appropriate ethics review.

### Chronic recordings

All the data used in this paper is described in Ref.^14^. Here, we focused on a subset of this data, i.e. the 18 mice which were chronically implanted with Neuropixels 2.0. During the experiments, mice were typically head fixed and exposed to sensory stimuli, engaged in a task, or resting. The mice were recorded from different experimental rigs, implanted, and recorded by different experimenters using different devices.

### DeepUnitMatch algorithm

The architecture of DeepUnitMatch reuses most of the architecture from UnitMatch, described in Ref.^14^. The main difference is its module to extract waveform feature similarities.

### Inputs to DeepUnitMatch

The first preprocessing step is to extract, for each neuron *i*, two features: a ‘snippet’ of its average spike waveform, and its position on the probe. The snippet corresponds to a 30-channel window of the average spike waveform of each neuron, centered on the channel where the voltage signal reaches its maximum (Figure 1e). For each neuron, two such snippets are extracted, corresponding to the average waveforms for each half of a recording. Thus, we define *w*_*c,t,r,i*_ the element of the snippet of neuron *i* and repeat *r*, at time *t* and on channel *c* . The position of each neuron **p**_*i*_ is defined as the spatial position of this maximum channel on the probe (2 dimensions).

### Network architecture

The DeepUnitMatch waveform similarity network is a convolutional network (Figure 1f), which takes individual waveform snippets (with channels from the two electrode columns interleaved) as an input. First, the snippet is passed through two convolutional blocks, one along the time dimension and one along the channel (spatial) dimension, while preserving its size. It is then passed through a spatial convolutional layer, which reduces its size to 15 x 60. Finally, it goes through three fully connected layers, which successively reduce its size to 512 and 256, its final size. The output of the network for neuron *i* and repeat *r* is then given by the vector **y**_*i,r*_ (of length 256 elements).

### Network training

Training of the neural network takes place in two steps, both of which are important to learn useful representations of the waveforms (Figure 1g,h and Supplementary Figure 2).

#### Training step 1: autoencoder

The first step is to extract general features of the neurons’ waveforms. We optimize the neural network to build the reconstructed waveform 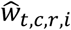 from the original input waveform *w*_*t,c,r,i*_, following the loss function *L*_*a*_:

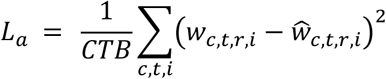

where *C* is the number of channels (30), *T* is the number of timepoints (60), *B* is the batch size, and *r* is chosen randomly for each neuron. This is a standard self-supervised autoencoder training framework^23^, which does not require any labels, and each snippet (and their repetition) is treated independently. The point of this step is to learn an informative compressed representation of the waveforms.

#### Training step 2: contrastive learning

The second step is to fine-tune the encoder to learn features that specifically differentiate different neurons. We use contrastive learning^24^, which is a powerful framework for unsupervised settings. A multi-layer perceptron with one hidden layer is used as a projection head to further compress the learned representations to 128-dimensional vectors. The cross-repeat cosine similarity *c*_*i,j*_ between the representation of neurons *i* and *j* is computed as follows:

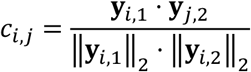

The neural network is trained in batches of size *B* using the Adam optimizer^47^. For each batch, we construct a *B* × *B* similarity matrix *S*, which contains similarity scores after row-wise SoftMax scaling. The entries *s*_*ij*_ of *S* are thus:

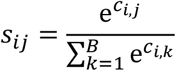

In this matrix, the diagonal elements correspond to scaled similarity scores between representations of copies of waveforms from the same neuron, while off-diagonal elements correspond to similarity scores between representations of waveforms from different neurons. Therefore, the goal of the fine-tuning step is to push this matrix towards an identity matrix. Only neurons from the same recording session are present in every batch.

We first define a weight matrix *W* which has diagonal elements *W*_*ii*_ = 1 and off-diagonal elements *W*_*ij*_ = 10. The cross-entropy loss is then applied to the Hadamard (entry-wise) product *S*^′^ = *S* ⊙ *W*. This incentivizes the network to push apart representations of waveforms from different units, thus optimizing it for the task at hand as opposed to learning generic compressed waveform representations. The final loss used for finetuning is:

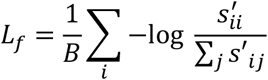

We show that while both steps help performance, the contrastive learning step is more important (Supplementary Figure 2). Moreover, in contrastive learning, augmentations are often used to reflect variations in the data which the neural network should learn to be invariant to. By having slightly varying copies of each waveform from the two halves, the training procedure naturally includes realistic augmentations, and we found that adding further transformations such as adding noise or spatial translation did not improve performance.

### Matching algorithm

After a pass through the trained network, each pair of waveforms is assigned a similarity score, which is the cosine similarity of their representations. The matching algorithm then uses this similarity score to track individual neurons across recordings, in several steps.

#### Merging split units (optional)

First, we use these scores to merge potential ‘split units’ within sessions, similarly to Ref.^19^. Split units occur when the upstream spike sorting algorithm that assigns spikes to neurons within a session erroneously splits a single neuron into two different units. We detected potential split units as pairs of different neurons from the same session that have waveform similarity above a certain threshold. The method for thresholding similarity is discussed below in the training of the naive Bayes model. We determine whether they can be merged using a model for the expected number of refractory period violations, given the number of spikes from a neuron in a certain recording^48^. If merging their spike trains results in fewer refractory period violations than expected, the two units are merged. We then create a new unit, the waveform of which is the average of the two merged units (weighted by the respective number of spikes) and its spike train formed by merging the two spike trains. We then re-trained the network on the data where split units have been merged and performed testing on this data.

#### Finding putative matches (1^st^ pass)

Next, we identify putative matches by applying a series of filters to the set of all possible pairs of neurons in a pair of sessions. The first filter thresholds the waveform similarity score at the crossing point of the distributions of *s*_*ii*_ and *s*_*ij*_ (*i* ≠ *j*) within sessions. To account for the difference in similarity distributions that may occur across sessions, we aligned the median of the within-session and across-session distributions. We further refine the resulting set of putative matches by directional and conflict filters. The directional filter ensures that both similarity values (*s*_*ij*_ and *s*_*ji*_ ) exceed the threshold for each match. The conflict filter ensures that no neuron is matched to more than one neuron in another session: only the highest-similarity match is kept. We thus obtain a matrix of matches *M* with elements *m*_*ij*_, where *m*_*ij*_ = 1 if the units are putative matches and 0 otherwise.

#### Correcting for drift

Using these putative matches, we corrected the position **p**_*i*_ of each neuron by removing the median difference of the position of all putatively matched pairs across sessions. We then computed the Euclidean distance matrix between pairs of neurons *E* with elements *e*_*ij*_:

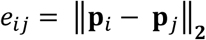

#### Finding putative matches (2^nd^ pass)

After drift correction, we recomputed putative matches, adding a spatial filter, which removes any potential matches that have a spatial distance greater than 20µm. The spatial filter was run after the directional filter and before the conflict filter.

#### Computing the match probabilities

Finally, we used the preliminary matches to build probability distributions for putative matching and non-matching pairs, for the waveform similarity scores *s*_*ij*_ and the drift-corrected distance between pairs. In this paper, to isolate the effect of our improvement of waveform similarity on performance, we reused the distance metric (drift-corrected centroid distance) provided by UnitMatch. Alternatively, in the final DeepUnitMatch pipeline, we use the drift-corrected Euclidean distance between pairs of units *e*_*ij*_. We then trained a Naive Bayes model with these distributions and putative matches labels to output the match probability for each pair of neurons *P*_*ij*_.

#### Finding final matches

To identify the final set of matches across pairs of sessions (Figure 2h,j and Figure 3a-d), we then selected the pairs with a match probability above 0.5, and applied the directional ( *P*_*ij*_ > 0.5 and *P*_*ji*_ > 0.5) and conflict (select bast matching pairs) filters. All pairs passing these criteria were considered as matches. We performed this step for both UnitMatch and DeepUnitMatch outputs.

#### Assigning unique identities

In Figure 3e,f, we used this probability matrix to assign a unique identification number for each neuron across recordings, as explained in Ref.^14^. We used the intermediate tracking algorithm from UnitMatch’s tracking module. To compute the probability of a unit being tracked, we then looked at each unit across all recordings and computed the probability of this unit being tracked in previous or subsequent recordings. These probabilities were then averaged across all the units from each animal, and averaged across animals.

#### Session exclusion criteria

AUCs values become highly noisy when there are too few neurons. In Figure 2h, only sessions where at least 20 matches were found by UnitMatch are included. In one mouse, UnitMatch couldn’t find any pair of sessions with at least 20 matches, and was thus excluded from this analysis. In Figure 2i,j and Figure 3a-d, only sessions where at least 20 matches were found by either UnitMatch or DeepUnitMatch are included.

### Model performance

We evaluated the model’s performance by computing a functional similarity score across pairs of neurons, using the inter-spike interval (ISI) histograms, as in Ref.^14^.

#### Functional similarity score

For each neuron and for each half of the recording, we compute the ISI histogram as the distribution of the times between consecutive spikes, binned on a logarithmic scale from 0 to 5 s. We then computed the functional similarity across pairs of neurons as the Pearson correlation coefficient between their ISI histograms.

#### AUC

Real matched pairs should have a higher similarity than real non-matched pairs. To test this, we computed the ROC curve for different populations of pairs: pairs coming from putative matched units, or nonmatched units, across days. We then computed the area under the ROC curve (AUC) to quantify this difference between distributions.

#### Simulations

To test the sensitivity of the AUC to false positives and false negatives (Supplementary Figure 1), we simulated experiment with 100 neurons over two days, with only 50 of these neurons being in both days. The functional similarity used to compute the AUC was defined as 1s for tracked neurons, 0 otherwise. We then computed the value of the AUC while varying the number of false positive or false negative errors.

## Supplementary figures

**Supplementary Figure 1.**
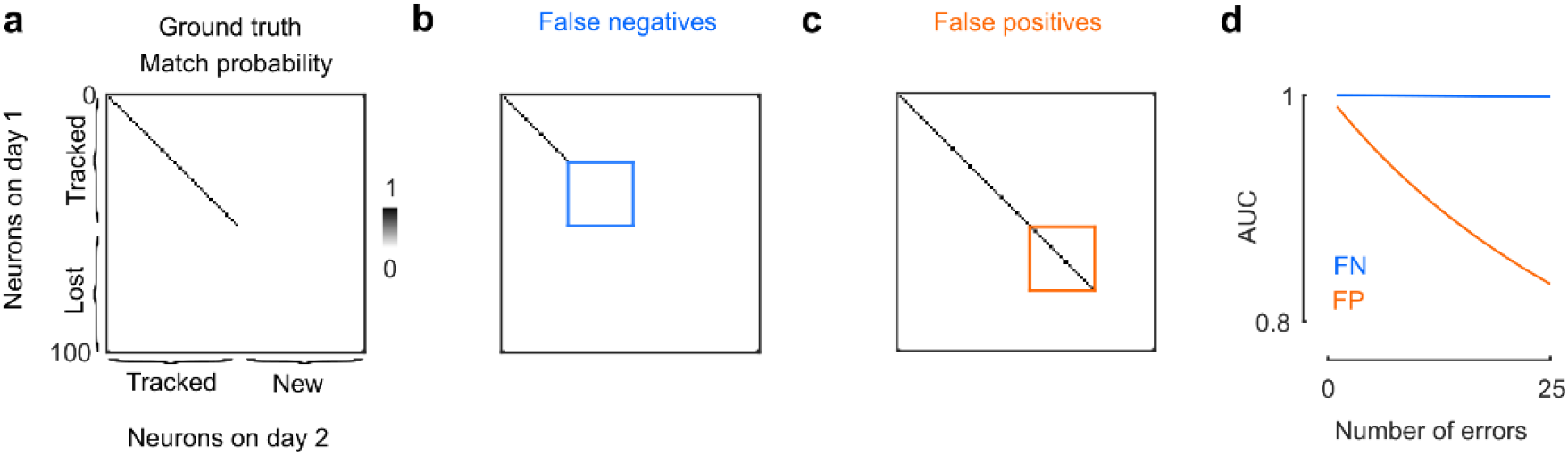
Sensitivity of our performance measure (AUC) to false positives and false negatives in a simulation. **a**, Simulated experiment with 100 neurons over two days, with only 50 of these neurons being in both days. In this extreme case, the functional similarity used to compute the AUC is defined as 1s for tracked neurons, 0 otherwise. **b**, Scenario where the matching algorithm has a high false negative rate. **c**, Scenario where the matching algorithm has a high false positive rate. **d**, Value of the AUC while varying the number of false positive (FP, *orange*) or false negative (FN, *light blue*) errors. Intuitively, this is because the number of potential matches (which will increase by 1 with a FP) scales with the number of neurons N, while the number of non-matches (which will increase by 1 with a FN) scales with N^2^.

**Supplementary Figure 2.**
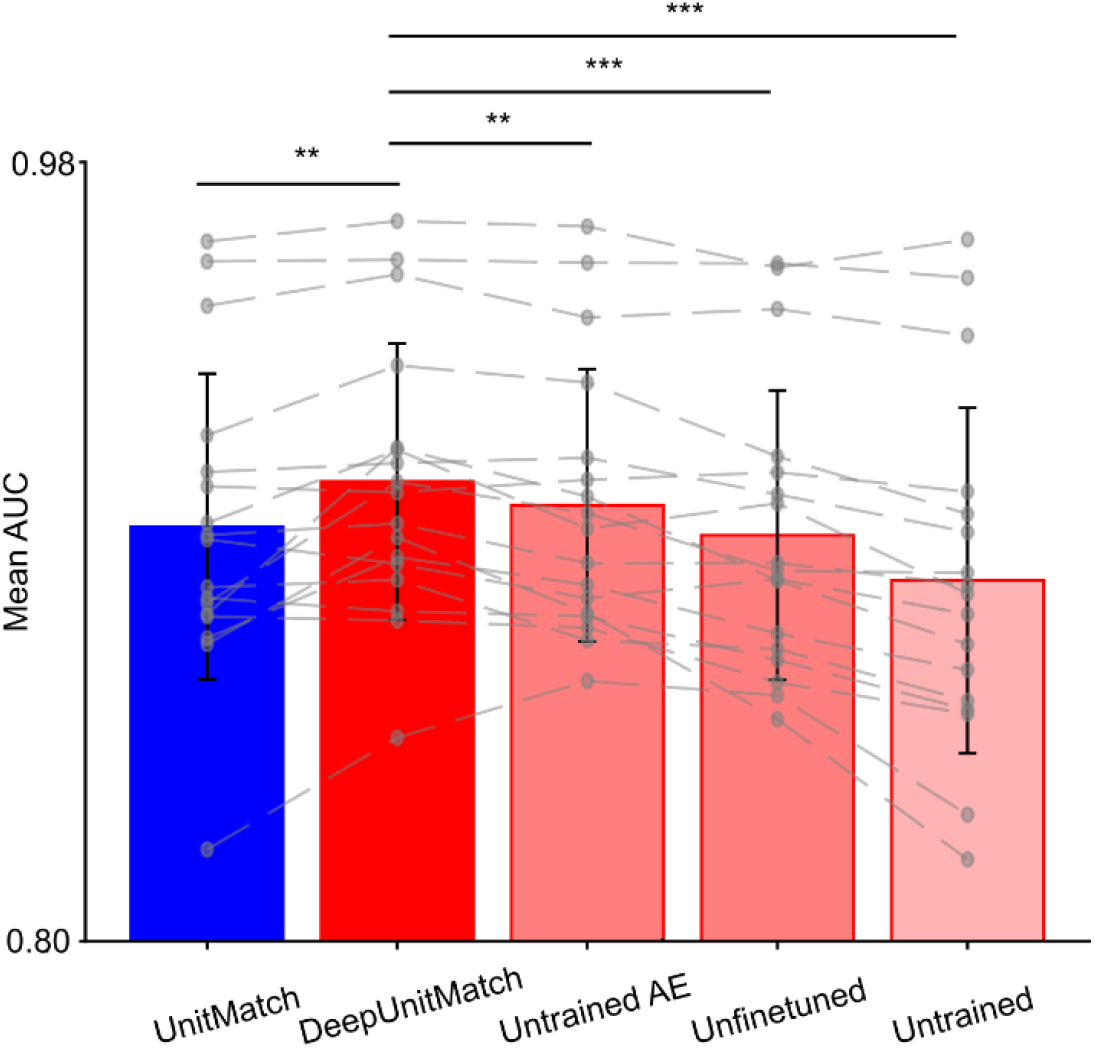
Performance of UnitMatch and DeepUnitMatch under different training configurations. Performance is quantified as the AUC averaged over mice, with the number of matches set by UnitMatch (*blue*). A minimum of 20 matches must be found for a pair of sessions to be included in these results. Bars indicate average ± s.t.d. AUC. Gray dashed lines for individual mice. **: p < 0.01, ***: p < 0.001

## Notes

### Competing Interest Statement

The authors have declared no competing interest.

